# Choanoflagellate transfection illuminates their cell biology and the ancestry of animal septins

**DOI:** 10.1101/343111

**Authors:** David S. Booth, Heather Szmidt-Middleton, Nicole King

## Abstract

As the closest living relatives of animals, choanoflagellates offer unique insights into animal origins and core mechanisms underlying animal cell biology. However, unlike traditional model organisms, such as yeast, flies and worms, choanoflagellates have been refractory to DNA delivery methods for expressing foreign genes. Here we report the establishment of a robust method for expressing transgenes in the choanoflagellate *Salpingoeca rosetta*, overcoming barriers that have previously hampered DNA delivery and expression. To demonstrate how this method accelerates the study of *S. rosetta* cell biology, we engineered a panel of fluorescent protein markers that illuminate key features of choanoflagellate cells. We then investigated the localization of choanoflagellate septins, a family of GTP-binding cytoskeletal proteins that are hypothesized to regulate the multicellular rosette development in *S. rosetta.* Fluorescently tagged septins localized to the basal pole of *S. rosetta* single cells and rosettes in a pattern resembling septin localization in animal epithelia. The establishment of transfection in *S. rosetta* and its application to the study of septins represent critical advances in the growth of *S. rosetta* as an experimental model for investigating choanoflagellate cell biology, core mechanisms underlying animal cell biology, and the origin of animals.

## INTRODUCTION

First described in the mid-19^th^ century, choanoflagellates inspired great debate regarding animal taxonomy (James-Clark, 1868; Kent, 1871; Leadbeater, 2015). The most diagnostic morphological feature of choanoflagellates, a “collar complex” composed of a single apical flagellum surrounded by a collar of actin-filled microvilli (Fig. 1), was interpreted as evidence of a special relationship between choanoflagellates and sponges, whose choanocytes (or “collar cells”) each bear a collar complex. Subsequent phylogenetic analyses and the discovery of cells with a collar complex in nearly all animal phyla have revealed that sponges and all other animals are monophyletic, with choanoflagellates as their closest living relatives (Fig. 1) (Lang *et al.*, 2002; King *et al.*, 2008; Ruiz-Trillo *et al.*, 2008; Burger *et al.*, 2003; Fairclough *et al.*, 2013; Brunet and King, 2017). Moreover, comparative genomic analyses have revealed that choanoflagellates, animals, and other holozoans express genes required for animal multicellularity and embryogenesis (Richter *et al.*, 2018; Sebe-Pedros *et al.*, 2017), including cadherins (Abedin and King, 2008), tyrosine kinases (Suga *et al.*, 2012; Manning *et al.*, 2008), and Myc (Young *et al.*, 2011). Thus, comparisons among animals and choanoflagellates have the potential to provide unique insights into animal origins and core features of animal cell biology that are not conserved in other experimental models, such as yeast (King, 2004).

**Figure 1:**
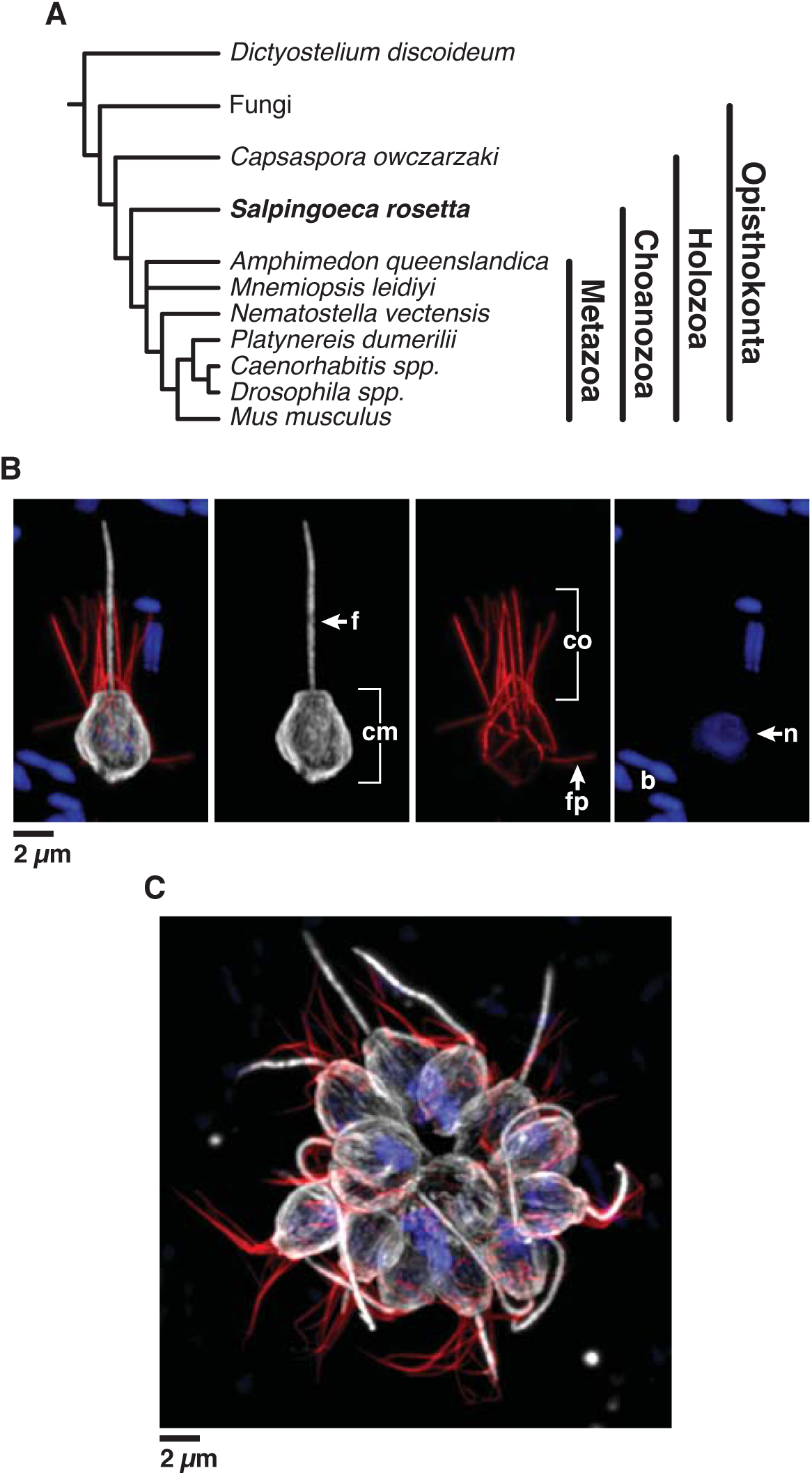
Introduction to *Salpingoeca rosetta*, an experimentally-tractable model choanoflagellate. **(A)** *S. rosetta* and other choanoflagellates (abbreviated as ‘choano’) are the closest living relatives of animals. **(B, C)** *S. rosetta* has a complex life history that includes single cells **(B)** and multicellular rosettes **(C)**. Immunofluorescence in fixed, permeabilized *S. rosetta* single cells **(B)** highlights the diagnostic cellular architecture of choanoflagellates, including a single apical flagellum (f) made of microtubules (white) surrounded by a collar (co) filled with F-actin (red) of microvilli. Staining for tubulin also illuminates cortical microtubules (cm) that run in parallel tracks along the cell periphery from the apical to the basal poles of each cell. DNA staining (blue) highlights the choanoflagellate nucleus (n) and the nucleoids of bacterial prey (b) present in choanoflagellate cultures. **(C)** In multicellular rosettes (stained as in panel **B**), the basal poles of cells are oriented toward the interior of the rosette and the apical flagella point outward.

The choanoflagellate *Salpingoeca rosetta* (previously named *Proterospongia* sp. (King *et al.*, 2003)) has recently emerged as an experimentally tractable model. *S. rosetta* develops from a single founding cell into a spherical, multicellular “rosette” (Fig. 1C) through serial rounds of cell division in a process that evokes the earliest stages of animal embryogenesis (Fairclough *et al.*, 2010). Since the establishment of the first *S. rosetta* cultures almost twenty years ago, *S. rosetta* has become increasingly amenable to cell and molecular biological approaches due to the sequencing of its genome (Fairclough *et al.*, 2013), the establishment of forward genetic screens (Levin and King, 2013; Levin *et al.*, 2014), the ability to experimentally control key events in its life history (Dayel *et al.*, 2011), and the discovery that environmental bacteria induce multicellular rosette development and mating (Alegado *et al.*, 2012; Dayel *et al.*, 2011; Woznica *et al.*, 2016; Woznica *et al.*, 2017).

An important remaining barrier to the study of molecular and cellular mechanisms in *S. rosetta* has been the inability to perform transfection and transgene expression. Furthermore, the absence of the RNA interference pathway in *S. rosetta* has precluded gene knockdowns (Richter *et al.*, 2018; Fairclough *et al.*, 2013). Here we report the establishment of a robust nucleofection-based method to transfect and express transgenes in *S. rosetta.* By engineering plasmids with *S. rosetta* regulatory sequences driving the expression of fluorescently-tagged *S. rosetta* proteins, we have developed a broad panel of markers for the study of choanoflagellate cell biology *in vivo*. As a first application, we used transgene expression to characterize septins, genes with conserved roles in fungal (Helfer and Gladfelter, 2006; Berepiki and Read, 2013) and animal development (Fairclough *et al.*, 2013; Adam *et al.*, 2000; Neufeld and Rubin, 1994; O’Neill and Clark, 2013; Kim, S. K. *et al.*, 2010), that have been hypothesized to regulate rosette development (Fairclough *et al.*, 2013). By imaging fluorescently-tagged septins in live cells, we show that their localization in *S. rosetta* resembles that in animal epithelia, providing a potential evolutionary link between the mechanisms underlying animal and choanoflagellate multicellularity.

## RESULTS

### A robust method to transfect *S. rosetta*

To detect successful transfection, we started by engineering four different DNA plasmid constructs, each with different *S. rosetta* regulatory sequences fused to a gene, *nanoluc* (Hall *et al.*, 2012), encoding a highly sensitive luciferase (Fig. S1B). Because no choanoflagellate promoters had previously been mapped, we increased the likelihood of cloning sequences that would drive robust gene expression by fusing *nanoluc* to non-coding sequences flanking a set of genes – *elongation factor L (efl), α-tubulin (tub), non-muscle actin (act)*, and *histone H3 (H3)* – that each exhibit high expression, lack introns in their open reading frames and have well-annotated 5’- and 3’-untranslated regions (Fig. S1A) (Fairclough *et al.*, 2013).

Next, we set out to deliver these DNA plasmid constructs into *S. rosetta* cells using nucleofection, an electroporation-based technique that has proven particularly effective for transfection of diverse eukaryotes (Janse, Ramesar *et al.*, 2006; Caro *et al.*, 2012; Vinayak *et al.*, 2015), including mammalian primary cells that are resistant to transfection (Gresch *et al.*, 2004; Hamm *et al.*, 2002). To quantify transfection efficiency, we performed luciferase assays on cell lysates. Through many trials, we eventually achieved a low level of transfection with nucleofection by improving conditions for culturing *S. rosetta* cells (Fig. S2), modifying approaches for handling cells throughout the nucleofection procedure (Supplementary Information), and screening thirty unique combinations of electrical pulses and buffers (Fig. S3).

Optimization around these initial conditions culminated in a procedure that provided robust and reproducible transfection of *S. rosetta* (Fig. 2A; Methods and http://www.protocols.io/groups/king-lab). When used in the optimized transfection procedure, all four transfection reporters drove strong expression of nanoluc protein, producing luminescence signals that were over three orders of magnitude above the detection limit (Fig. 2B).

**Figure 2:**
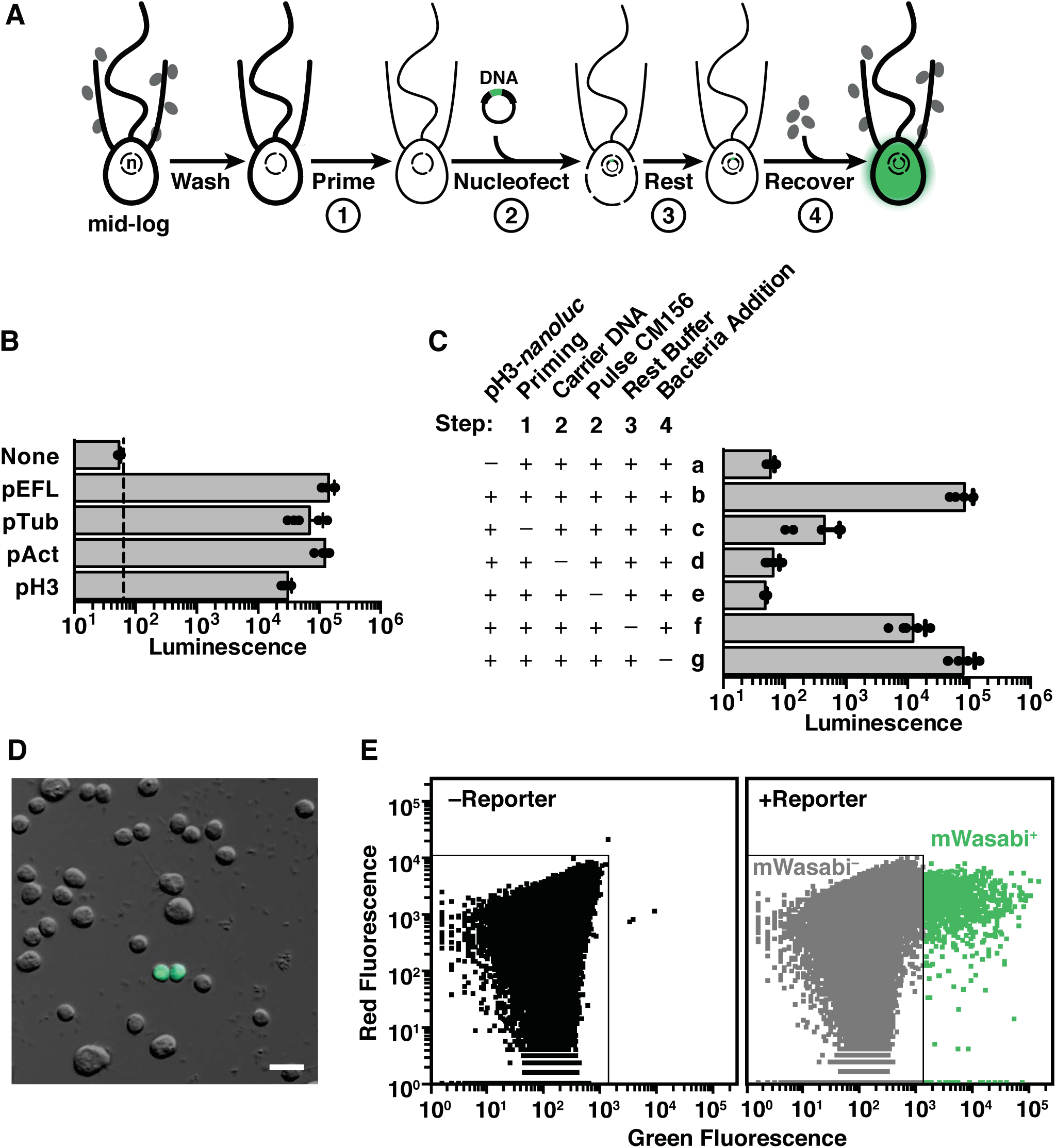
A robust procedure for transfecting *S. rosetta*. **(A)** A summary of the step-wise procedure to transfect *S. rosetta* with DNA plasmids. To prepare *S. rosetta* for transfection, cells were harvested at mid-log phase and then washed to remove bacteria (depicted as grey ovals). *S. rosetta* cells (depicted with an apical collar, flagellum, and nucleus; n) were primed for nucleofection (step 1) through washing with a buffer that degrades extracellular material. A DNA plasmid encoding a highly sensitive luciferase, nanoluc, or a fluorescent protein was then transfected into the nucleus with a nucleofector (step 2). Immediately after transfection, the cells rested in a buffer that promotes membrane closure (step 3). Finally, the cells were transferred into 1x High Nutrient Media prepared with AK seawater for two days (step 4) before we assayed the expression of nanoluc or fluorescent proteins from the transfected DNA. **(B)** Non-coding DNA sequences flanking the coding sequences for *S. rosetta elongation factor L* (pEFL), *α-tubulin* (pTub), *β-actin* (pAct), and *histone H3* (pH3) genes drive the expression of a codon-optimized *nanoluc* reporter gene. 2.5 *µ*g of pEFL-*nanoluc,* pTub-*nanoluc,* pAct-*nanoluc*, and pH3-*nanoluc* reporter plasmids were each transfected into *S. rosetta* and the cells were subsequently assayed for luciferase expression. Each reporter produced a luminescence signal that was at least three orders of magnitude greater than the detection limit (dotted line) and significantly greater (one-way ANOVA, *p* < 0.001) than the background from a negative control, in which cells were transfected with an empty pUC19 vector (None). See Materials and Methods for details on replicates and statistical tests. **(C)** Systematically omitting each step of the transfection procedure revealed critical steps for the delivery and expression of plasmid DNA in *S. rosetta* cells. As a baseline for comparison, cells with 2.5 *µ*g of pH3-*nanoluc* reporter (row b) produced a luciferase signal that was three orders of magnitude greater than the background detected from cells transfected without the reporter plasmid (row a). Omitting the priming step by incubating cells in artificial seawater instead of priming buffer (row c), decreased luciferase signal by over two orders of magnitude. Nucleofection without carrier DNA (row d) or the application of the CM156 electrical pulse (row e) resulted in a complete loss of luciferase signal, indicating that both were essential for successful transfection. Directly transferring cells to sea water after nucleofection instead of a buffer that promotes membrane resealing during the rest step (row f) decreased the luciferase signal almost ten-fold. Finally, despite the fact that most prey bacteria were washed out prior to nucleofection, addition of fresh prey bacteria did not appear to be necessary. Supplementing transfected cells with fresh prey bacteria at the start of the recovery step had seemingly little effect on transfection success (row g), probably due to the persistence of a small number of live bacteria throughout the nucleofection procedure. **(D and E)** Fluorescent reporters mark transfected cells. Live cells transfected with a pAct-m*Wasabi* reporter construct could be observed by fluorescence microscopy **(D)** and quantified by flow cytometry **(E)**. Untransfected cells were used to draw a gate that includes 99.99% of cells, or four-standard deviations above the mean fluorescence value (left). The same gate was applied to a population of transfected cells (right) to categorize the mWasabi-population. Cells with higher values of green fluorescence that laid outside of the mWasabi-gate are categorized as mWasabi+. The efficiency of transformation, as quantified by three independent flow cytometry experiments, was ∼1% in a population of 1 million cells.

Because this was the first example, to our knowledge, of successful transgene expression in any choanoflagellate, we sought to identify which steps in the optimized protocol were most essential. Using the p*H3*-*nanoluc* transfection reporter, we quantified how the omission of each step impacted transfection efficiency (Fig. 2C). In addition to the use of an optimal electrical pulse during nucleofection, the two most important steps were priming the cells through the enzymatic removal of the extracellular matrix prior to nucleofection (Fig. 2A, step 1; Fig. S4) and the inclusion of carrier DNA during nucleofection (Fig. 2A, step 2; Fig. S5); eliminating either of these steps resulted in a nearly complete loss of signal.

Priming the cells for nucleofection was a novel step motivated by our observation that *S. rosetta* cells are surrounded by a potentially protective extracellular coat (Dayel *et al.*, 2011; Levin *et al.*, 2014; Leadbeater, 2015), and the inclusion of carrier DNA (pUC19) in nucleofection reactions eliminated the need to include large quantities of reporter construct plasmid (Fig. S5). Another improvement was the development of a recovery buffer that enhanced transfection ten-fold (Fig. 2A, step3), presumably by promoting membrane resealing after nucleofection (Rols and Teissie, 1990; Rols and Teissie, 1989).

Luciferase assays performed on cell lysates gave a sensitive read-out of population-wide nanoluc expression but did not allow the examination of live, transfected cells nor reveal the proportion of cells that were successfully transfected. Therefore, we next engineered eight reporters with different fluorescent proteins placed under the control of regulatory sequences from the *S. rosetta actin* homolog. Fluorescence was readily detected from cells transfected with reporters encoding mTFP1 (Ai *et al.*, 2006), mWasabi (Ai *et al.*, 2008), sfGFP (Pedelacq *et al.*, 2006), mNeonGreen (Shaner *et al.*, 2013), mPapaya (Hoi *et al.*, 2013), TagRFP-T(Shaner *et al.*, 2008), mCherry (Shaner *et al.*, 2004), and tdTomato (Shaner *et al.*, 2004). In contrast, an eGFP (Yang *et al.*, 1996) reporter failed to yield fluorescent cells, likely due to protein misfolding, as cells transfected with the ‘super-folder’ variant of GFP (sfGFP) did fluoresce properly. In transfected cells, the fluorescent signal was distributed throughout the nucleus and cytosol yet excluded from membrane bound compartments (Fig. 3A).

**Figure 3:**
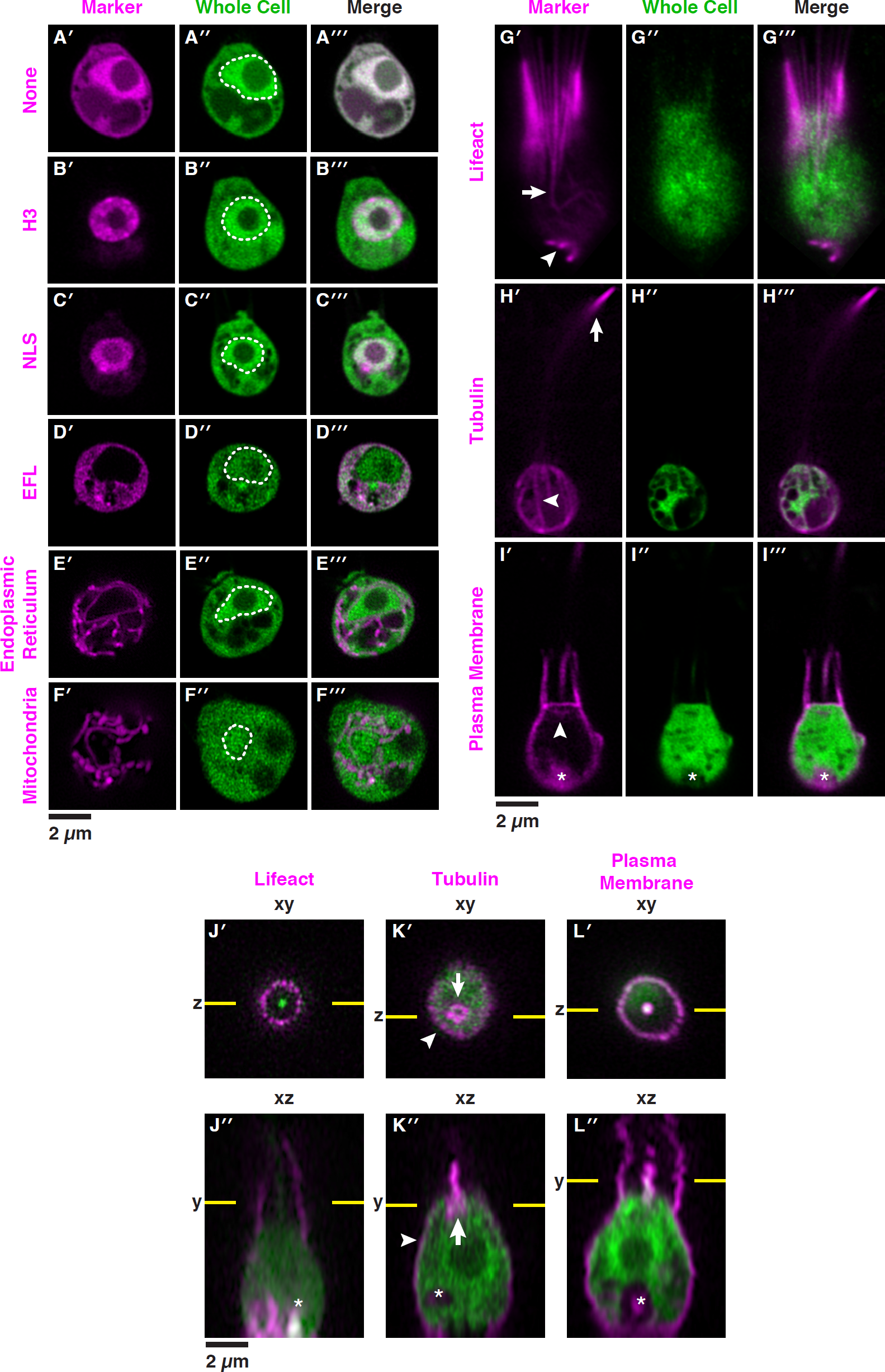
Fluorescent markers illuminate the cell biology of *S. rosetta* in live cells. Fluorescent subcellular markers expressed from reporter plasmids in live *S. rosetta* cells were constructed by fusing *mCherry* in frame to genes encoding localization peptides and proteins (Datasets S1 and S3). Twenty-four hours after co-transfecting cells with 5 *µ*g of a plasmid encoding a subcellular marker fused to the mCherry protein and 5 *µ*g of a plasmid encoding untagged mTFP1 that served as a whole cell marker, live cells were visualized by superresolution microscopy with a Zeiss LSM 880 Airyscan. The variation in localization of the whole cell mTFP1 marker stems from cell-to-cell differences in the number and localization of vacuoles, which exclude mTFP1. In panels **A** -**I**, the cells are oriented with the apical flagellum at the top and the nucleus, when included in the plane of focus (**A’’ - F’’)**, is indicated with a dotted white line. **(A)** Without localization signals (None), fluorescent proteins (mCherry, **A’**, and mTFP1, **A’’**) were distributed throughout the cell with a slight enrichment in the nucleus and complete exclusion from other membrane bound compartments. **(B and C)** A fusion of mCherry to the carboxy terminus of Histone H3 **(B’)** or the amino terminus of a simian virus 40 nuclear localization signal (NLS; **C’)** was confined to the nucleus, whereas mCherry fused to the carboxy terminus of elongation factor L (EFL; **D)** was excluded from the nucleus and restricted to the cytosol. **(E)** The endoplasmic reticulum (ER) was highlighted by fusing the signal sequence from Rosetteless (PTSG_03555) and an ER retention sequence (HDEL from PTSG_07223) to the amino and carboxy termini of mCherry, respectively. **(F)** The mitochondrial network was highlighted by fusing a targeting sequence from *S. cerevisiae* CoxIV to the amino terminus of mCherry. **(G)** A Lifeact peptide fused to the amino terminus of mCherry marked filamentous actin (F-actin) that forms filipodia (arrowhead) and actin filaments in the cell body that coalesce to form the collar (arrow). **(H)** Fusing mCherry to the amino terminus of α-Tubulin highlighted parallel tracks of microtubules (arrowhead) that extended subcortically from the apical pole to the basal pole of cells and microtubules that emerged from the apical pole of the cell body to form the flagellum. The flagellum undulates rapidly in live cells and can be difficult to image in total; in this cell the most distal tip of the flagellum is captured in the plane of focus (arrow). **(I)** A plasma membrane marker constructed by fusing a geranyl-geranylation sequence (PTSG_00306) to the carboxy terminus of mCherry outlined the entire cell shape, including the collar, flagellum, and cell body. The membrane marker also weakly highlighted the Golgi (arrowhead). The food vacuole (asterisk) was often visualized due to autofluorescence from ingested bacteria or through accumulation of the fluorescent markers in the food vacuole, perhaps through autophagy. **(J - L)** Orthogonal views along the xy and xz axes from confocal micrographs showed fine details of cell architecture that were highlighted by transfecting cells with F-actin, microtubule, and plasma membrane markers fused to mCherry (magenta). In xz views, each cell is oriented with the flagellum facing toward the top of the micrograph; the flagella appeared shorter and blurred because of the sigmoidal shape of the flagellar beat. Lifeact **(J)** and the plasma membrane **(L)** markers fused to mCherry showed the microvilli (arrowheads). **(K)** The α-tubulin-mCherry showed the subcortical tracks of microtubules at the cell periphery (arrowhead) and the microtubule organizing center (arrow).

We observed that transfected cells resembled untransfected cells in their shape, motility, and ability to propagate, indicating that transfection did not irreparably harm *S. rosetta*. Fluorescence persisted through multiple cell divisions, yet the diminishing signal in daughter cells indicated that transfection was transient (Fig. S6A). Importantly, using flow cytometry one to two days after transfection, we found that ∼1% of the population was reproducibly transfected, and fluorescence-activated cell sorting enriched this transfected cell population (Fig. S6B). This transfection frequency is comparable to high frequency episomal transformation of the model yeast *Saccharomyces cerevisiae* that ranges from 1 – 10% (Schiestl and Gietz, 1989; Kawai *et al.*, 2010), and similar transfection frequencies are achieved in model apicocomplexans (Janse, Franke-Fayard *et al.*, 2006; Caro *et al.*, 2012).

### Fluorescent markers illuminate the cell architecture of *S. rosetta*

To demonstrate the versatility of the new method for transfection and simultaneously explore the cell biology of *S. rosetta in vivo*, we designed a set of fluorescent reporters to mark key features of *S. rosetta* cells: the nucleus, cytoplasm, collar, filopodia, flagellum, membrane, mitochondria and endoplasmic reticulum (ER). For each fluorescent reporter, the *mCherry* gene was fused in-frame to *S. rosetta* DNA sequences encoding conserved proteins or peptides that localize to specific organelles or subcellular regions in yeast and mammalian cells. To benchmark each fluorescent marker, we compared its localization in transfected cells to cellular landmarks known from electron and immunofluorescence micrographs (Fig. S7) (Abedin and King, 2008; King *et al.*, 2009; King *et al.*, 2008; Sebe-Pedros *et al.*, 2013; Leadbeater, 2015).

Electron micrographs have revealed two distinct regions in the nucleus: the darkly-stained nucleolus positioned in the center and the surrounding, more lightly-stained nucleoplasm (Leadbeater, 2015; Burkhardt *et al.*, 2014). As predicted, mCherry fused to either the carboxy terminus of H3 or the amino terminus of the simian virus 40 nuclear localization signal localized primarily to the *S. rosetta* nucleoplasm and was excluded from the cytoplasm (Fig. 3B, C) (Kanda *et al.*, 1998; Kalderon *et al.*, 1984). In contrast, the cytoplasmic marker EFL-mCherry (Huh *et al.*, 2003) localized to the cytosol and was excluded from the nucleus (Fig. 3D).

Two of the most diagnostic features of the choanoflagellate cell are the actin-filled collar and the flagellum, which is comprised of microtubules (Karpov and Leadbeater, 1998). A fusion of mCherry to the filamentous actin-binding peptide Lifeact (Riedl *et al.*, 2008) highlighted the parallel arrangement of straight microvilli in the collar (Fig. 3G and 3J), as well as filopodia extending from the basal pole of the cell (Fig. 3G, lower arrow)(Karpov and Leadbeater, 1998; Sebe-Pedros *et al.*, 2013). In live cells, Lifeact-mCherry revealed the native structure of the collar, which can be distorted in cells fixed for staining with fluorescent phalloidin or actin antibodies (Sebe-Pedros *et al.*, 2013). Lifeact-mCherry also showed details of actin filament organization that have not previously been evident, such as the existence of actin filaments that originate in the cell body and coalesce at the base of the collar to form each microvillus (Fig. 3G’, upper arrow; improved immunofluorescence techniques also preserve these cortical actin filaments, Fig. 1B). A fusion of α-tubulin to mCherry (Straight *et al.*, 1997) illuminated individual cortical microtubules emanating from the base of the flagellum to the basal pole of the cell (Fig. 3K’, arrow) and allowed visualization of the rapidly beating flagellum in live cells (Fig. 3H and 3K).

A cell membrane marker, with a geranyl-geranylation sequence fused to mCherry (Wang and Casey, 2016; Reid *et al.*, 2004), outlined the entire cell, including the flagellum, collar, and cell body (Fig. 3I and 3L), and faintly marked the Golgi apparatus (Fig. 3I’, arrow). In live cells, the cell membrane marker captured the formation of a phagocytic cup engulfing bacterial prey (Fig. S8)(Dayel and King, 2014). The ER marker (Friedman *et al.*, 2011), which included the amino terminal signal sequence from the secreted protein Rosetteless (Levin *et al.*, 2014) and a carboxy terminal ER retention sequence from the ER resident chaperone BiP (PTSG_07223), highlighted the continuity of the ER with the nuclear envelope and the distribution of ER throughout the cell, including around vacuoles (Fig. 3F). A mitochondrial marker (Friedman *et al.*, 2011) with an amino terminal targeting sequence from *S. cerevisiae* Cytochrome C Oxidase, Subunit IV revealed a network of mitochondria (Nunnari *et al.*, 1997) that is enriched around the nucleus and extends throughout the cell (Fig. 3G). Taken together, these fluorescent markers demonstrate new experimental capabilities to rapidly tag proteins and to monitor their localization in distinct cellular compartments and locales.

### Transgenesis reveals septin localization in live *S. rosetta* single cells and rosettes

A major motivation for establishing transgenics in *S. rosetta* was to rapidly characterize candidate genes for multicellularity. Therefore, we investigated the localization of septins, a family of paralogous genes hypothesized to contribute to multicellular development in *S. rosetta* (Fairclough *et al.*, 2013). The assembly of septin monomers into higher order structures is important for septin localization (McMurray *et al.*, 2011) and for the conserved roles of septins in regulating cytokinesis and cell polarity in fungi and animals. In animals, septins also function in phagocytosis (Huang *et al.*, 2008), ciliogenesis (Hu *et al.*, 2010), and planar cell polarity (Kim *et al.*, 2010). Septin proteins have a characteristic domain architecture with a diagnostic amino terminal guanosine triphosphate binding domain (G-domain), and most septins also have a carboxy terminal coiled-coil domain (Pan *et al.*, 2007; Nishihama *et al.*, 2011) (Fig. 4A). Septin paralogs interact directly through their G-domains to form heteromeric filaments, and these heteromeric filaments interact with each other through the septin coiled-coil domains to form higher order assemblies (Bertin *et al.*, 2008; Garcia *et al.*, 2011; Sirajuddin *et al.*, 2007).

**Figure 4:**
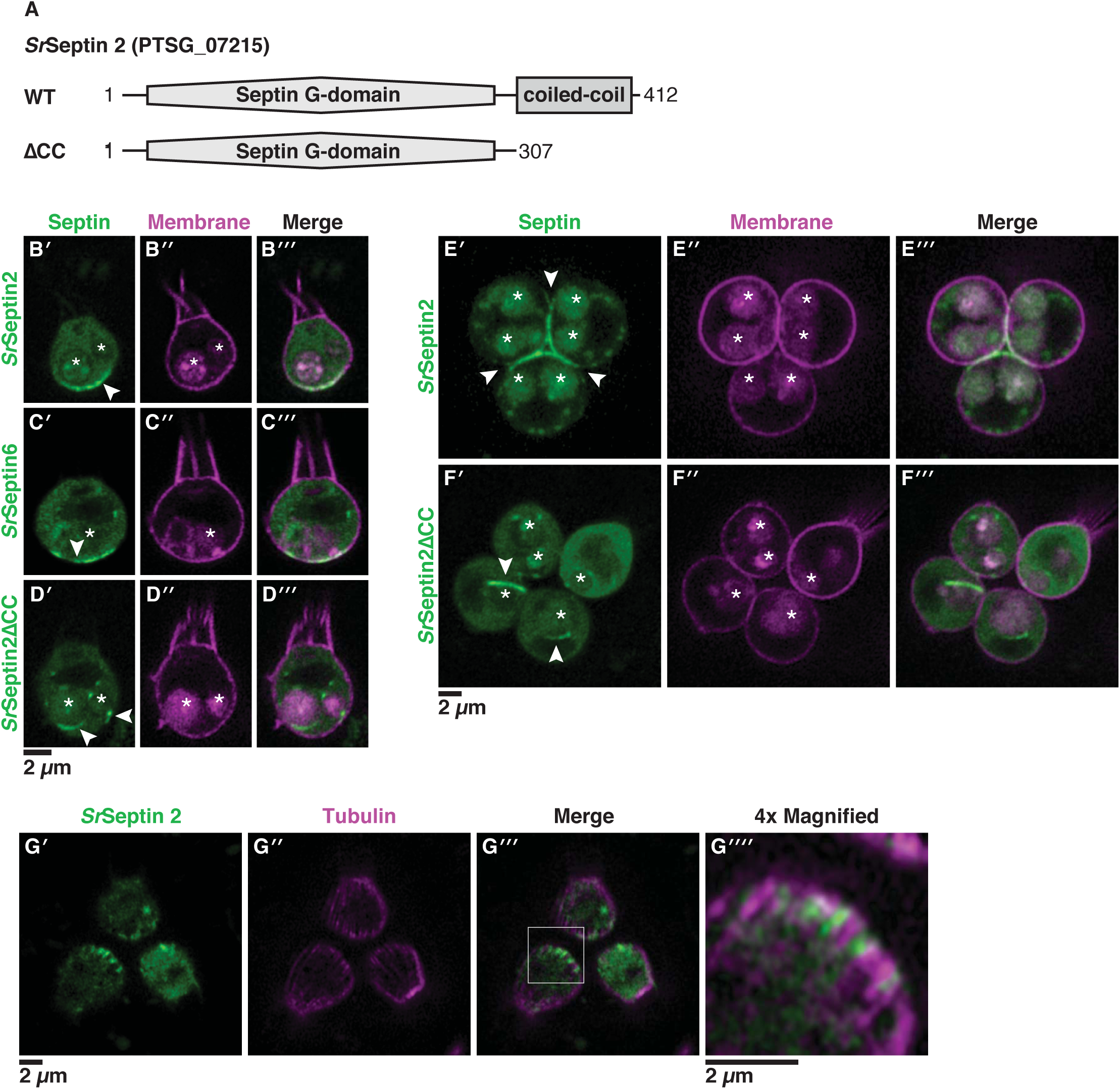
Septins assemble at the basal pole of *S. rosetta* cells. **(A)** *Sr*Septin2 has a prototypical protein domain architecture of septins, with an amino-terminal Septin G-domain that mediates filament formation and a carboxy terminal coiled-coil domain that mediates higher order assemblies of septin filaments. To investigate the localization of *Sr*Septin2, we engineered fusions with mTFP1 at the amino terminus and created a truncation of the coiled-coil domain (ΔCC). **(B)** A mTFP1-*Sr*Septin2 fusion protein localized to the basal pole of unicellular cells (**B’**, arrowhead). Co-transfecting cells with mTFP1-*Sr*Septin2 and a plasma membrane marker revealed *Sr*Septin2 distributed throughout the cytosol and enriched at the basal pole in confocal slices through the center of the cell. **(C)** mTFP1-*Sr*Septin6 mirrored the enrichment of mTFP1-*Sr*Septin2 at the basal pole (**C’**, arrowhead). The overlapping localization of *Sr*Septin2 and *Sr*Septin6 was compatible with these proteins forming heteromeric filaments with each other and other septin paralogs. **(D)** Consistent with the coiled-coil domain mediating the localization of septins through the formation of higher-order structures, *Sr*Septin2ΔCC localized throughout the cytoplasm, with no visible enrichment at the basal pole. Surprisingly, the deletion also caused ectopic filaments (**D’**; arrowheads) to form around membrane-bound vesicles that were, based on their size and position in the cell, presumably food vacuoles. **(E)** In rosettes, mTFP1-*Sr*Septin2 localized to points of cell-cell contact corresponding to the basal poles of cells (**E’**; arrowhead). **(F)** As in single cells, mTFP1-*Sr*Septin2ΔCC in rosettes was distributed throughout the cytosol and formed ectopic filaments (**F’**; arrowheads) around vacuoles. In panels **E** and **F**, *S. rosetta* single cells were transfected as in panels **B** and **C**, immediately induced to develop into rosettes (Woznica *et al.*, 2016), and imaged the next day. **(G)** *Sr*Septin2 intercalated between microtubules at the basal pole of the cell. Co-transfecting cells with mTFP1-*Sr*Septin2 and the α-tubulin marker showed *Sr*Septin2 filaments intercalated between microtubules at the basal pole in confocal slices that capture the cell cortex to easily visualize microtubule tracks. (**G’**, **G’’**, **G’’’**; box). **G’’’’** shows a 4x magnification of the basal pole of a representative cell (boxed region from **G’**, **G’’**, **G’’’**). In panels B-F, autofluorescence from ingested bacteria or through accumulation of the fluorescent markers highlights the food vacuole (asterisk).

*S. rosetta* expresses four septin paralogs, three of which have well-annotated genome models in the current draft genome (Fairclough *et al.*, 2013). We started by examining the localization of the *S. rosetta* septin protein *Sr*Septin2 (PTSG_07215), a septin with domains conserved in human and fungal septins (Fairclough *et al.*, 2013) that are necessary for septin filament formation (Sirajuddin *et al.*, 2007; Bertin *et al.*, 2008). Strikingly, mTFP1-*Sr*Septin2 was enriched at the basal poles of single and rosettes cells (Fig. 4B and 4E) and at points of contact between adjacent cells in rosettes (Fig. 4E). This mTFP1-*Sr*Septin2 fusion likely revealed the native localization of *Sr*Septin2 in *S. rosetta* because septins visualized by immunofluorescence microscopy in yeast (Ford and Pringle, 1991; Kim, H. B. *et al.*, 1991; Cid *et al.*, 1998; Haarer and Pringle, 1987), *Drosophila* (Adam *et al.*, 2000; Silverman-Gavrila *et al.*, 2008; Neufeld and Rubin, 1994), and mammalian cells (Spiliotis *et al.*, 2008) display the same localization as septins tagged with fluorescent proteins.

To investigate whether *Sr*Septin2 localized to the basal pole of cells might be part of heteromeric septin filaments, we examined the localization of another septin paralog, *Sr*Septin6 (PTSG_06009) and found that *Sr*Septin6 displays the same basal localization as *Sr*Septin2 (Fig. 4C). Such colocalization (Fig. S9) and the sequence homology with septins that have previously been shown to form heteromeric filaments (Sirajuddin *et al.*, 2007) strongly suggests that *Sr*Septin2 and *Sr*Septin6 assemble together at the basal pole. We further found that the basal localization of *Sr*Septin2 requires the coiled-coil domain, as a complete deletion of the coiled-coil domain (SrSeptin2ΔCC; Fig. 4A, D, F) eliminated *Sr*Septin2 enrichment at the basal pole when expressed in wild-type cells. Unexpectedly, mTFP1-*Sr*Septin2ΔCC formed ectopic rings around vesicles in the cytosol in wild-type cells (Fig. 4D and 4F). The localization of mTFP1-*Sr*Septin2ΔCC resembled the formation of ectopic septin filaments at convex membranes and the depletion of septins at the concave membranes in the filamentous fungi *Ashbya gosypii* (Meseroll *et al.*, 2012) upon the deletion of the coiled-coil domain of the fungal septin Shs1p. Similarly, vesicles to which mTFP1-*Sr*Septin2ΔCC localized have convex membranes as opposed to the concave membrane at the basal end of *S. rosetta* where wild-type *Sr*Septin2 localized.

The basal and lateral localization of *Sr*Septin2 and *Sr*Septin6 in rosettes is reminiscent of septin localization in polarized epithelial cells (Fares *et al.*, 1995; Spiliotis *et al.*, 2008), in which septins interact with the positive ends of microtubules that are growing toward the basal pole (Bowen *et al.*, 2011). In choanoflagellates, microtubules radiate down from the apical microtubule organizing centers (Karpov and Leadbeater, 1998), with the plus ends meeting at the basal pole of each cell, similar to the orientation of microtubule plus ends toward the basal pole in animal epithelia (Meads and Schroer, 1995). To examine if septins also interact with the plus ends of microtubules in *S. rosetta*, we co-transfected cells with mTFP1-*Sr*Septin2 and the tubulin marker α-tubulin-mCherry (Fig. 4G). Fluorescence microscopy showed that septin filaments intercalate between cortical microtubules at the basal pole of the cell (Fig. 4G). These data are consistent with conserved interactions between septins and microtubules from yeast to animals (Kusch *et al.*, 2002; Kremer *et al.*, 2005; Sellin *et al.*, 2011; Spiliotis *et al.*, 2008), including at the plus-ends of microtubules in choanoflagellates and animal epithelia (Bowen *et al.*, 2011).

## DISCUSSION

By synthesizing our growing knowledge of *S. rosetta* biology with a rigorous characterization and optimization of each step in the transfection procedure, we have developed a robust method for transgenesis in *S. rosetta* that can be easily implemented by other laboratories. This method overcomes numerous barriers that prevented efficient DNA delivery in our prior attempts using diverse methods, including standard electroporation, lipofection, bombardment, and cell-penetrating peptides. A key breakthrough for this study was the discovery that the extracellular coat surrounding *S. rosetta* might present a barrier for transfection, which motivated the development of a method to gently remove the extracellular material surrounding *S. rosetta,* thereby sensitizing cells for transfection. Additional improvements to the transfection procedure, such as a step for promoting the closure of the plasma membrane after electrical pulsation, were designed to address the unique challenges that arise from culturing *S. rosetta* in sea water. Just as our method was informed by approaches developed in model microeukaryotes (*Chlamydomonas* and yeast), the methods we have established in *S. rosetta* will likely extend to aid gene delivery in diverse non-model marine microeukaryotes. Overall, the gestalt of continually improving choanoflagellate husbandry (Levin and King, 2013), developing protocols for priming and recovering cells during nucleofection, and extensively optimizing transfection based on a quantitative assay produced a robust method for gene delivery in *S. rosetta*.

This work also provides a foundational set of vectors for expressing transgenes in *S. rosetta* (Dataset S1). In these vectors, the expression of luciferase or fluorescent proteins was placed under the control of native regulatory elements. From these vectors, we constructed a panel of fluorescently tagged subcellular markers that serve as references for monitoring the localization of other proteins in *S. rosetta*. For example, through our pilot study of *Sr*Septin2 and *Sr*Septin6, the use of these new transgenic tools revealed that septins localize to the basal pole of choanoflagellates, mirroring their localization in animal epithelial cells (Spiliotis *et al.*, 2008; Fares *et al.*, 1995).

Observing septin localization in *S. rosetta* contributes to our understanding of how septin functions evolved prior to the evolution of an epithelium in stem animals. A hallmark of animal epithelia is tightly connected cells that each have an apical and basal pole comprised of distinct lipids, proteins, and cytoskeletal structures (Rodriguez-Boulan and Macara, 2014). Septins help shape epithelia and other types of cells by serving as buttresses (Tanaka-Takiguchi *et al.*, 2009; Tooley *et al.*, 2009) and diffusion barriers (Barral *et al.*, 2000; Takizawa *et al.*, 2000; Hu *et al.*, 2010) for membranes that have distinct geometries (Bridges *et al.*, 2016; Cannon *et al.*, 2018) and lipid compositions (Casamayor and Snyder, 2003; Tanaka-Takiguchi *et al.*, 2009; Bertin *et al.*, 2010; Bridges *et al.*, 2014) and by interacting with microtubules (Spiliotis, 2010) and filamentous actin (Mavrakis *et al.*, 2014). While comparisons between animals and fungi have revealed conserved functions of septins in cell organization (Spiliotis and Gladfelter, 2012), the divergence of the lineages that gave rise to fungi and animals over one billion years ago (Parfrey *et al.*, 2011) resulted in important differences in fungal and animal cell biology (Stajich *et al.*, 2009). In fungi, septins facilitate polarized cell growth toward the new daughter cell (Kim *et al.*, 1991; Berepiki and Read, 2013), compartmentalize connected daughter cells (Barral *et al.*, 2000; Takizawa *et al.*, 2000; Helfer and Gladfelter, 2006), and mediate cytokinesis (Hartwell, 1971). In addition to their conserved roles in cell division (Neufeld and Rubin, 1994), animal septins have specific functions in epithelia that maintain apical-basal polarity (Spiliotis *et al.*, 2008), planar cell polarity (Kim *et al.*, 2010), intercellular adhesion (Kim, J. and Cooper, 2018; Park *et al.*, 2015), and ciliogenesis (Hu *et al.*, 2010; Kim *et al.*, 2010). The basal localization of septins in *S. rosetta* suggests that septin filaments organize a distinct region at the basal end of the cell, perhaps supporting intercellular contacts at the basal ends of cells in rosettes (Fig. 4E). Consistent with this hypothesis is the observation that the Rosetteless protein, which is necessary for rosette development, localizes to the basal end of cells prior to secretion into the interior of rosettes where the basal ends of cells meet (Levin *et al.*, 2014). Continued study of septin function in *S. rosetta* will establish the mechanisms by which septins facilitate multicellular development, and further comparisons with non-metazoan holozoans with recently established transgenic methods, such as *Capsaspora owczarzaki* (Parra-Acero *et al.*, 2018) and *Creolimax fragrantissima* (Suga and Ruiz-Trillo, 2013), will provide further insights into the ancestral functions of septins.

Previous analyses of gene function in choanoflagellates relied on custom antibodies (Abedin and King, 2008; Young *et al.*, 2011; Burkhardt *et al.*, 2014; Levin *et al.*, 2014), laborious forward genetic screens (Levin *et al.*, 2014), and *in vitro* biochemistry (Burkhardt *et al.*, 2014). The ability to express transgenes in *S. rosetta* described here will accelerate studies of the ancestral functions of animal genes that are conserved in choanoflagellates. We anticipate that future work will build on this approach, eventually leading to the development of methods for stable transgenesis and genome editing in *S. rosetta*. Combining an expanded repertoire of approaches for investigating gene function in-depth in *S. rosetta* with comparisons to other experimentally-tractable choanoflagellates (Li *et al.*, 2018; Richter *et al.*, 2018) and non-choanozoans (Suga and Ruiz-Trillo, 2013; Parra-Acero *et al.*, 2018) promises to yield increasingly mechanistic insights into the ancestry of animal cell biology.

## MATERIALS AND METHODS

### Cell culture and media preparation

*S. rosetta* was cultured with a single bacterial species, *E. pacifica*, that serves as a food source (Levin and King, 2013). Media recipes are provided in Table S1. Cultures were established from frozen aliquots by adding 1 ml of thawed cells to 10 ml of 0.2x High Nutrient Media (Table S1). After the cells reached a density of 10^4^ cells/ml, the culture was split 1:2 into 1x High Nutrient Media with a constant volume of 0.24 ml/cm^2^. After this initial split (denoted as day 0), cells were passaged in 1x High Nutrient Media according to the following schedule: 1:4 dilution on day 1, 1:8 dilution on day 2, 1:16 on day 3. Subsequently cells were passaged every day at a 1:24 dilution or every other day as a 1:48 dilution of cells.

Based on the recommendation from Lonza to use a medium with a low calcium concentration for transfecting mammalian cells, we searched for a seawater recipe with a lower concentration of calcium than the routinely-used artificial seawater made from Tropic of Marin Salts (Levin and King, 2013), which has a calcium concentration of 9.1 mM at a salinity of 35 g/kg (Atkinson and Bingman, 1998). The AK seawater formulation (Table S1) that has been used to culture marine algae (Hallegraeff *et al.*, 2004) and dinoflagellates (Skelton *et al.*, 2009) and has a calcium concentration of 2.7 mM. We found that *S. rosetta* grows more rapidly in 1x High Nutrient Media prepared in AK seawater rather than seawater prepared with Tropic of Marin Salts (Fig. S2A). Therefore, we switched to a growth medium based on AK seawater for routine culturing. After optimizing the nucleofection protocol, we demonstrated that growing *S. rosetta* in AK seawater also resulted in higher transfection efficiencies (Fig. S2B) than growing *S. rosetta* in seawater prepared with Tropic of Marin Salts.

### Reporter plasmid design and molecular cloning

Dataset S1 lists the complete inventory of engineered plasmids with a summary of primers, cloning methods, and annotations for constructing each plasmid. Complete plasmid sequences and plasmids have also been deposited at Addgene (http://www.addgene.org/Nicole_King). Below is a brief summary of considerations for designing plasmids, and a more detailed description of standard molecular cloning methods for engineering plasmids can be found in the Supplementary Information.

#### Cloning regulatory regions from *S. rosetta* genes

Because we had no previous knowledge about the architecture of choanoflagellate regulatory regions, we aimed to clone as much as 1000 bp upstream and downstream of targeted open reading frames as these fragments are slightly larger than the mean intergenic distance of 885 bp (Sebe-Pedros *et al.*, 2017). Of necessity, the cloned intergenic sequences reported here were shorter to avoid repetitive CA and GT sequences that were present before the putative promoter and after the 3’-UTR, respectively. To increase the specificity of primers, we designed the primers to anneal to regions with a GC content ≤ 50%, as the *S. rosetta* genome is 56% GC. Ultimately, the cloned regions that encompass the promoter and the 5’-UTR ranged in size from 550 bp to 1095 bp and those encompassing the 3’UTR ranged from 200 bp to 807 bp.

#### Synthetic gene design

Synthetic reporter genes (*nanoluc* and the genes encoding diverse fluorescent proteins listed in Supplementary File 1) were codon optimized to match the codon usage of the set of highly expressed intron-less genes listed in Fig. S1, as codon usage can be biased for highly expressed genes (Hiraoka *et al.*, 2009). A codon usage table (Dataset S2) was generated from the coding sequences of highly expressed intronless genes (Dataset S2A) and from all coding sequences (Dataset S2B) using the ‘cusp’ tool in Emboss (Rice *et al.*, 2000). The codon usage table was then used to generate a codon optimized DNA sequence for each target protein sequence with the ‘backtranseq’ tool on Emboss. The DNA sequences were further edited by making synonymous substitutions with less frequently used codons to change restriction enzyme sites and to remove repetitive sequences. Finally, sequences were added to the ends of these designed genes for cloning with restriction enzymes or Gibson assembly. The engineered reporter gene sequences are available through Addgene (Dataset S1; http://www.addgene.org/Nicole_King).

#### Subcellular marker design

Dataset S3 provides the amino acid sequences for all of the subcellular markers reported in Fig. 3. To ensure that the fluorescent protein tag for each marker would not interfere with the functions of proteins or peptides that determine localization, some of the constructs were engineered to have a flexible linker sequence (SGGSGGS) separating the fluorescent protein and the localization signals.

### Optimized transfection protocol

The protocol is summarized in Fig. 2 and detailed protocols for reagent preparation and transfection are available at protocols.io at the following link: http://www.protocols.io/groups/king-lab

#### Culture

Two days prior to transfection, a culture flask (Corning, Cat. No. 353144) was seeded with *S. rosetta* at a density of 5,000 cells/ml in 200 ml of 1x High Nutrient Media. The culture was supplemented with 2 mg of frozen *E. pacifica* by resuspending a 10 mg pellet of flash-frozen *E. pacifica* in 1 ml of media and then adding 200 *µ*l of the resuspended pellet to the culture of *S. rosetta*.

#### Wash

After 36-48 hours of growth, bacteria were washed away from *S. rosetta* cells through three consecutive rounds of centrifugation and resuspension in sterile AK seawater. The culture flask was vigorously shaken for 30 sec to homogenize the 200 ml that was seeded two days prior (see above) and then transferred to 50 ml conical tubes and spun for 5 min at 2000 x g and 22°C. The supernatant was removed with a serological pipette, and residual media was removed with a fine tip transfer pipette. The cell pellets were resuspended in a total volume of 100 ml of AK seawater, vigorously shaken in their conical tubes for 30 sec, and then centrifuged for 5 min at 2200 x g and 22°C. The supernatant was removed as before. Each cell pellet was resuspended in 50 ml of AK seawater, vigorously shaken for 30 sec, and centrifuged for 5 min at 2400 x g and 22°C. After the supernatant was removed, the cells were resuspended in a total volume of 100 *µ*l of AK seawater. A 100-fold dilution of cells was counted on a Luna-FL automated cell counter (Logos Biosystems) and the remaining cells were diluted to a final concentration of 5×10^7^ choanoflagellate cells/ml. The resuspended cells were divided into 100 *µ*l aliquots with 5×10^6^ cells per aliquot to immediately prime cells in the next step. A 200 ml culture typically yields 6-8 aliquots of cells.

#### Prime

After washing away bacteria, each aliquot of *S. rosetta* cells was incubated in priming buffer to remove the extracellular material coating the cell. The 100 *µ*l aliquots that contained 5×10^6^ cells were centrifuged for 5 min at 800 x g and at room temperature. The supernatant was removed with a fine tip micropipette. Cells were resuspended in 100 *µ*l of priming buffer (40 mM HEPES-KOH, pH 7.5; 34 mM Lithium Citrate; 50 mM L-Cysteine; 15% (w/v) PEG 8000; and 1 *µ*M papain) and then incubated for 30 min. Priming was quenched by adding 2 *µ*l of 50 mg/ml bovine serum albumin-fraction V (Sigma) and then centrifuged for 5 min at 1250 x g and 22°C with the centrifuge brake set to a ‘soft’ setting. The supernatant was removed with a fine-tip micropipette, and the cells were resuspended in 25 *µ*l of SF Buffer (Lonza).

#### Nucleofect

Each transfection reaction was prepared by adding 2 *µ*l of ‘primed’ cells resuspended in SF buffer to a mixture of 14 *µ*l of SF Buffer; 2 *µ*l of 20 *µ*g/*µ*l pUC19; 1 *µ*l of 250 mM ATP, pH 7.5; 1 *µ*l of 100 mg/ml Sodium Heparin; and ≤7 *µ*l of reporter DNA. (Note that higher volumes of nucleofection lead to lower transfection frequencies; thus, reporter DNA should be as concentrated as possible, not exceeding 7 *µ*l. Also, see below for “Note about titrating reporter plasmids.”) The transfection reaction was transferred to one well of a 96-well nucleofection plate (Lonza). The nucleofection plate was placed in a Nucleofector 4d 96-well Nucleofection unit (Lonza), and the CM156 pulse was applied to each well.

#### Rest and recover

Immediately after pulsation, 100 *µ*l of ice-cold recovery buffer (10 mM HEPES-KOH, pH 7.5; 0.9 M Sorbitol; 8% (w/v) PEG 8000) was added to the cells, Recovery buffer was gently mixed with the transfected cells by firmly tapping the side of the plate and then incubating the samples for 5 min. The whole volume of the transfection reaction plus the recovery buffer was transferred to 1 ml of 1x High Nutrient Media in a 12-well plate. After the cells recovered for 1 hour, 5 *µ*l of a 10 mg frozen *E. pacifica* pellet resuspended in media (see above), was added to each well. The cells were grown for 24 to 48 hours before assaying for luminescence or fluorescence.

#### Note about establishing transfection in non-model microeukaryotes

Establishing a transfection protocol for *S. rosetta* required adapting several different transfection procedures for a variety of eukaryotic cells to meet the unique requirements of *S. rosetta*. While the specific details for transfecting *S. rosetta* may not be readily applicable to other organisms, the general considerations and the process for optimization that led to the development of the transfection protocol described here could inform efforts to transfect other microeukaryotes. Therefore, we have included a summary in the supplementary information (See text and Figs. S10 and S11) of the initial development and optimization of the aforementioned protocol.

### Nanoluc reporter assay

To measure relative transfection efficiency resulting from different transfection protocols and promoters, we performed luciferase assays on lysates of transfected cells. Cells transfected with 2.5 *µ*g of *nanoluc* reporter plasmids were pelleted by centrifuging for 10 min at 4200 x g and 4°C. The supernatant was removed and the cells were resuspended in 50 *µ*l of NanoGlo buffer (Promega) and then transferred to a well of a white, opaque 96-well plate (Greiner Bio-one Cat No.655083). Luminescence was immediately recorded on a Spectramax L Microplate Reader (Molecular Devices) with a 1 min dark adaption and 10 sec dwell time with the photomultiplier gain set to photon counting mode.

Based on standard definitions from analytical chemistry (Harris, 2007) the detection limit was set to three standard deviations above the background signal such that any signal above the detection limit has less than a 1% chance of arising from random error. The limit of detection was calculated in two different ways. First, the y-axis intercept and standard deviation were calculated from a standard curve (Harris, 2007) fit to a serial dilution of nanoluc versus luciferase activity (Fig. S3A). To decrease the bias toward higher luciferase values, the standard curve was fit with the objective 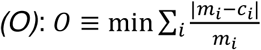, where *m* is the measured luciferase value for a given data point *i* and *c* is the calculated luciferase value. Second, the detection limit was also determined as three standard deviations above the mean of eight replicate luciferase measurements of cells transfected without any reporter plasmid, which resulted in the same calculated detection limit.

Reproducibility in luciferase assays was assessed by performing at least two independent experiments on separate days with different preparations of ‘primed’ cells; data presented in Fig. 2, S2, S3, S10, and S11 represent one of the independent experiments. Within each experiment from the same preparation of ‘primed’ cells, replicate measurements were performed by setting up three to five independent transfections for each condition (shown as black dots); bar graphs in Fig. 2, S2, S3, S10, and S11 show the mean values of the five independent transfections with error bars showing the standard deviation. Before performing statistical tests that rely on a normal distribution, luciferase data were transformed to a log-normal distribution by taking the base-10 logarithm of luciferase values as gene expression data from luciferase assays display a log-normal distribution (Muranaka *et al.*, 2013).

### Flow Cytometry

To measure the percentage of cells expressing each of the different transgenes under different transfection conditions, we used flow cytometry. Cells were transfected with 10 *µ*g of mWasabi or 10 *µ*g of TagRFP-T reporter plasmids for flow cytometry because these fluorophores produced the highest fluorescence signal upon illumination with the 488 or 561 nm lasers, respectively. To prepare cells for flow cytometry, cultures from 10-12 transfections were pooled 24 hours after transfection and centrifuged for 15 min at 3600 x g and 4°C. The supernatant was removed with a fine-tip transfer pipette to avoid disturbing the pellet. The pelleted cells were resuspended in 500 *µ*l of 0.22 *µ*m filtered AK seawater and then filtered through a 40 *µ*m filter.

Because a large number of bacteria were present in the cultures, *S. rosetta* cells were gated based on the area of forward scattering signal versus the area of the side scattering signal and the area of the forward scattering signal versus the height of the forward scattering signals. To differentiate transfected cells from untransfected cells, fluorescence signal was measured using lasers and filters for the fluorophores FITC (Green Fluorescence) and PE (Red Fluorescence); untransfected cells form a population along the y=x line of these plots, and the population of transfected cells are skewed along one axis that corresponds to the fluorophore. The transfected cells were gated to exclude >99.99% of untransfected cells as determined from a negative control reaction that was transfected without a fluorescent reporter (Fig. 2E, left panel).

### Live cell imaging

An important benefit of transgenics is the ability to visualize protein localization and cell architecture in living cells. To this end, we have established improved protocols for live cell imaging in *S. rosetta*. Glass-bottomed dishes were prepared for live cell microscopy by corona-treating the glass for 10 s. Afterwards, 300 *µ*l of 0.1 mg/ml poly-D-lysine was applied to the glass cover (18 *µ*l/cm^2^), incubated for 10 min at room temperature, and then removed. Excess poly-D-lysine was washed away from the glass surface with three rinses of 500 *µ*l artificial seawater.

Cells transfected with 5 *µ*g of each fluorescent reporter were prepared for microscopy by centrifuging 1-2 ml of transfected cells for 10 min at 3,600 x g and 4°C. After centrifugation, the supernatant was removed and the cell pellet was resuspended in 200 *µ*l of 4/5 Tropic of Marin artificial seawater with 100 mM LiCl. Lithium chloride slows flagellar beating, as in spermatozoa (Brokaw, 1987; Gibbons and Gibbons, 2013), to decrease the movement of cells during imaging. The resuspended cells were pipetted on top of the poly-D-lysine coated glass-bottom dish and adsorbed on the surface for 10 min. Lastly, 200 *µ*l of 20% (w/v) Ficoll 400 dissolved in 4/5 Tropic of Marin artificial seawater with 100 mM LiCl was pipetted drop-by-drop on top of the cells. The addition of Ficoll decreases flagellar movement by increasing the viscosity of the media (Wilson *et al.*, 2015; Pate and Brokaw, 1980) without significantly changing the osmolarity or refractive index of the sample (GE Lifesciences).

Confocal microscopy was performed on a Zeiss Axio Observer LSM 880 with an Airyscan detector and a 63x/NA1.40 Plan-Apochromatic oil immersion objective. The mTFP1 and mCherry fluorophores were selected for two-color imaging due to their high photostability and minimal spectral overlap. Confocal stacks were acquired in superresolution mode using ILEX line scanning and two-fold averaging and the following settings: 40 nm x 40 nm pixel size, 93 nm z-step, 0.9-1.0 *µ*sec/pixel dwell time, 850 gain, 458 nm laser operating at 5% laser power, 561 nm laser operating at 3% laser power, 458/561 nm multiple beam splitter, and 495-550 nm band-pass/570 nm long-pass filter. Images were initially processed using the automated Airyscan algorithm (Zeiss) and then reprocessed by setting the Airyscan threshold 0.5 units higher than the value reported from automated Airyscan processing. The stacks were further processed by correcting for signal decay, background, and flickr in Zen Blue (Zeiss). Last, FIJI (Schindelin *et al.*, 2012) was used to apply a gamma factor to each channel and subtract the background using a 100 pixel radius.

Epifluorescence and differential interference contrast images were recorded using a Zeiss Axio Observer.Z1/7 Widefield microscope with a Hamamatsu Orca-Flash 4.0 LT CMOS Digital Camera and 40x/NA 1.1 LD C-Apochromatic water immersion, 63x/NA1.40 Plan-Apochromatic oil immersion, or 100x NA 1.40 Plan-Apochromatic oil immersion objectives. Green fluorescent proteins were imaged with a 38 HE filter set and red fluorescent proteins with a 43 HE filter set. Images were processed by applying a gamma factor and background subtracting fluorescence channels in FIJI.

#### Note about titrating reporter plasmids

A titration of fluorescent reporter plasmids showed that 10 *µ*g of total reporter plasmid(s) best balanced transfection efficiency, brightness, and a faithful indication of subcellular architecture. We caution that high plasmid concentrations can result in the overexpression of fluorescent markers, leading to aberrant localization of the marker and gross changes in cell morphology. Such artefacts can be avoided by performing a titration to determine the best concentration of plasmid and recording images from cells with a range of fluorescence intensities that result from any transfection. One of the best markers to assess optimal reporter plasmid concentrations is the tubulin marker because of its distinct localization that can be benchmarked with immunofluorescence.

### Immunofluorescence staining and imaging

Immunfluorescence was performed as previously described (Woznica *et al.*, 2016) with modifications to better preserve features of the cytoskeleton. Two milliliters of cells were concentrated by centrifugation for 10 min at 2750 x g and 4°C. The cells were resuspended in 400 *µ*l of artificial seawater and applied to poly-L-lysine coated coverslips (BD Biosciences) placed in the bottom of each well of a 24-well cell culture dish. After allowing the cells to settle on the coverslip for 30 min, 150 *µ*l of the cell solution was gently removed from the side of the dish. It is crucial to leave a small layer of buffer on top of cells to preserve the cell morphology, hence the 250 *µ*l of liquid left in the well. All of the subsequent washes and incubations during the staining procedure were performed by adding and removing 200 *µ*l of the indicated buffer.

Cells were fixed in two stages. First, the coverslip was washed once with 6% acetone in cytoskeleton buffer (10 mM MES, pH 6.1; 138 KCl, 3 mM MgCl_2_; 2 mM EGTA; 675 mM Sucrose), which better preserves the actin cytoskeleton (Cramer and Mitchison, 1995; Symons and Mitchison, 1991), and then incubated for 10 min at room temperature after a second application of the acetone solution. Subsequently, the coverslip was washed once with 4% formaldehyde diluted in cytoskeleton buffer and then incubated for 15 min at room temperature after a second application of the formaldehyde solution. Last, the coverslip was gently washed three times with cytoskeleton buffer.

Cells were permeabilized by washing the coverslip once with permeabilization buffer (100 mM PIPES, pH 6.95; 2 mM EGTA; 1 mM MgCl_2_; 1% (w/v) bovine serum albumin-fraction V; 0.3% (v/v Triton X-100) and then incubated for 30 min upon a second addition of permeabilization buffer. After the permeabilization buffer was removed, the coverslip was washed once with primary antibody, 50 ng/ml mouse E7 anti-tubulin antibody (Developmental Studies Hybridoma Bank) diluted in permeabilization buffer, and then incubated for 1 h in a second application of primary antibody. The coverslip was gently washed twice in permeabilization buffer. Next, the coverslip was washed once with secondary antibody, 8 ng/ml Donkey anti-mouse IgG–AlexaFluor568 (ThermoFisher) diluted in permeabilization buffer, and then incubated for 1 h after a second application of secondary antibody. Afterwards, the coverslip was washed once in permeabilization buffer and then three times with PEM (100 mM PIPES-KOH, pH 6.95; 2 mM EGTA; 1 mM MgCl_2_). The coverslip was washed once with 10 *µ*g/ml Hoechst 33342 and 4 U/ml Phalloidin-AlexaFluor488 in PEM and then incubated for 30 min with a second application of Hoechst33342/Phalloidin. Finally, the coverslip was washed once in PEM.

To prepare a slide for mounting, 10 *µ*l of Pro-Long Diamond (Invitrogen) was added to a slide. The coverslip was gently removed from the well with forceps, excess buffer was blotted from the side with a piece of filter paper, and the coverslip was gently placed on the drop of Pro-Long diamond. The mounting media was allowed to cure overnight before visualization.

Images were acquired on a Zeiss LSM 880 Airyscan confocal microscope with a 63x objective (as described for live cell imaging) by frame scanning in the superresolution mode with the following settings: 35 nm x 35 nm pixel size; 80 nm z-step; 0.64 *µ*s/pixel dwell time; 561 nm laser operating at 1.5% power with a 488/561 nm beam splitter, a 420-480 nm/495-620 nm band pass filter, and a gain of 750; 488 nm laser operating at 1.5% power with a 488/561 nm beam splitter, a 420-480 nm/495-550 nm band pass filter, and a gain of 750; and 405 nm laser operating at 1.5% power with a 405 nm beam splitter, a 420-480 nm/495-550 nm band pass filter, and a gain of 775.

## Acknowledgements

Laura Wetzel, Monika Sigg, Hannah Elzinga, Lily Helfrich, and Reef Aldayafleh helped with experiments and reagent preparation. Corey Allard, as part of the Marine Biological Laboratory’s Physiology Course, helped with early tests of priming conditions. Kent McDonald generously provided a transmission electron micrograph from samples prepared by Pawel Burkhardt. We thank these people for providing access and support for scientific instruments: Russell Vance and Lab for use of their luminometer, Hector Nolla and Alma Valeros in the Flow Cytometry Facility, and the UC Berkeley DNA Sequencing Facility. The following individuals generously donated reagents and provided technical support: Brad Hook and Dee Czarniecki from Promega, Ethan Brooks from Lonza, and Colleen Manning from Zeiss. We appreciate scientific discussions and advice from these individuals: David Schaffer, Sabrina Sun, Jorge Ortiz, Niren Murthy, Tara DeBoer, Fyodor Urnov, Matt Welch and Lab, Rebecca Heald and Lab, Abby Dernberg and Lab, and Amy Gladfelter. We thank members of the King lab for helpful discussions, research support, and comments on the manuscript, especially Arielle Woznica, Kayley Hake, and Ben Larson. An additional thanks to the following people for providing comments on the manuscript: Candace Britton, Pawel Burkhardt, Matt Daugherty, Galo Garcia, Tera Levin, Kristin Patrick, and Dan Richter. DSB is supported as a Simons Foundation Postdoctoral Fellow of the Jane Coffin Childs Memorial Fund for Biomedical Research. This work was funded in part by a grant from the Gordon and Betty Moore Foundation’s Marine Microbiology Initiative for establishing Emerging Model Systems.

## Author Contributions

DSB and NK conceived of the project and wrote the manuscript. DSB, HSM, and NK designed experiments and interpreted data. DSB and HSM collected data.

